# Reference ontology and database annotation of the COVID-19 Open Research Dataset (CORD-19)

**DOI:** 10.1101/2020.10.04.325266

**Authors:** Oliver Giles, Rachael Huntley, Anneli Karlsson, Jane Lomax, James Malone

**Author notes:** **Corresponding authors** Oliver Giles, James Malone.

## Abstract

The COVID-19 Open Research Dataset (CORD-19) was released in March 2020 to allow the machine learning and wider research community to develop techniques to answer scientific questions on COVID-19. The dataset consists of a large collection of scientific literature, including over 100,000 full text papers. Annotating training data to normalise variability in biological entities can improve the performance of downstream analysis and interpretation. To facilitate and enhance the use of the CORD-19 data in these applications, in late March 2020 we performed a comprehensive annotation process using named entity recognition tool, TERMite, along with a number of large reference ontologies and vocabularies including domains of genes, proteins, drugs and virus strains. The additional annotation has identified and tagged over 45 million entities within the corpus made up of 62,746 unique biomedical entities. The latest updated version of the annotated data, as well as older versions, is made openly available under GPL-2.0 License for the community to use at: https://github.com/SciBiteLabs/CORD19

## Introduction

In March 2020 in response to the COVID-19 pandemic, a consortium including The White House, AI2, CZI, MSR, Georgetown and NIH released an open research dataset along with a call for action [1] to the science and technology community. They called on artificial intelligence experts to develop new text and data mining techniques that can help the science community answer high-priority scientific questions related to COVID-19. The original dataset can be accessed at Kaggle: https://www.kaggle.com/allen-institute-for-ai/CORD-19-research-challenge

Data normalisation is an important step in the data preparation phase of many machine learning approaches [2]. The use of ontologies to normalise this data has shown improvements in various machine learning applications [3, 4]. Using ontologies to standardise descriptions of data also offers benefits in post-analysis. Using the same ontology identifiers in annotations can enable the integration of results across multiple disparate but similarly annotated datasets. The value of sharing well-annotated data has never been made clearer with the emergence of COVID-19 [5, 6] and it is this challenge we are addressing here.

To contribute to this call for action, in March 2020, and for the months following, we developed updated COVID-19 focused vocabularies replete with rich synonyms to identify relevant virus strain names, proteins and genes often studied with these viruses, and related entities such as gene functions and processes. These vocabularies use public identifiers where available from the likes of NCBI Taxon, Uniprot and the Gene Ontology. Using these vocabularies along with others describing indications, phenotypes, species, countries, genes, proteins and drugs, we applied our named entity recognition tool, TERMite, to annotate the entire CORD-19 dataset. This has produced a richly annotated dataset with over 45 million annotations, using 62,746 unique biomedically relevant entities. In order to facilitate the discovery of connections between entities in the corpus, we have produced a set of entity cooccurrences. This includes sentences which mention at least two entities, for instance a drug and gene. Because the vocabularies for identifying COVID-19 related terms are available, this can be used to filter for COVID-19 specific co-occurrences.

## Method

The CORD-19 dataset is described as “the most extensive machine-readable coronavirus literature collection available for data mining” [7]. As of September 2020, the dataset contains over 200,000 scientific articles, including over 100,000 with full text, focusing on COVID-19, SARS-CoV-2, and other relevant or related coronaviruses. The dataset is made available as a set of JSON documents, each document includes the title, authors, abstract, body text and detailed references (where appropriate).

To identify relevant concepts, a focused set of vocabularies was required with enrichments specific to concepts pertinent to COVID-19 research. The new COVID-19 specific vocabularies, and a set of existing vocabularies, mostly aligned to public ontologies (Table 1).

**Table 1.**
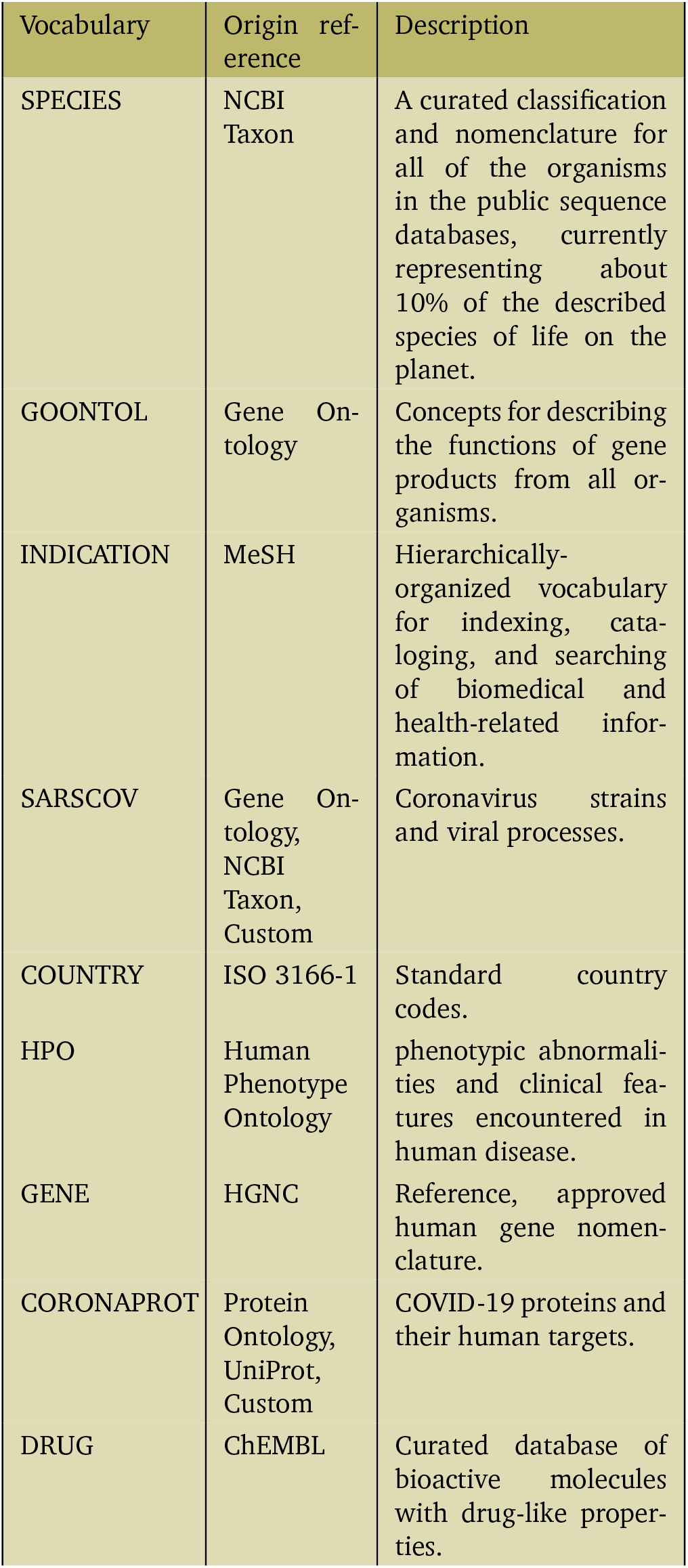
Summary of the vocabularies used to annotate the CORD-19 data in this work.

| Vocabulary | Origin reference | Description |
| --- | --- | --- |
| SPECIES | NCBI Taxon | A curated classification and nomenclature for all of the organisms in the public sequence databases, currently representing about 10% of the described species of life on the planet. |
| GOONTOL | Gene Ontology | Concepts for describing the functions of gene products from all organisms. |
| INDICATION | MeSH | Hierarchically-organized vocabulary for indexing, cataloging, and searching of biomedical and health-related information. |
| SARSCOV | Gene Ontology, NCBI Taxon, Custom | Coronavirus strains and viral processes. |
| COUNTRY | ISO 3166-1 | Standard country codes. |
| HPO | Human Phenotype Ontology | phenotypic abnormalities and clinical features encountered in human disease. |
| GENE | HGNC | Reference, approved human gene nomenclature. |
| CORONAPROT | Protein Ontology, UniProt, Custom | COVID-19 proteins and their human targets. |
| DRUG | ChEMBL | Curated database of bioactive molecules with drug-like properties. |

**Table 2.** Summary of the NER annotations across each vocab used.

| Vocabulary | Total hits | Unique hits |
| --- | --- | --- |
| SPECIES | 9958967 | 3502 |
| GOONTOL | 6975073 | 10702 |
| INDICATION | 12998639 | 6360 |
| SARSCOV | 2629924 | 302 |
| COUNTRY | 2078689 | 228 |
| HPO | 3997204 | 5737 |
| GENE | 4018135 | 15714 |
| CORONAPROT | 795869 | 67 |
| DRUG | 2513491 | 20134 |

To create the annotations for each document, a commercial named entity recognition engine (NER), TERMite, was employed using these vocabularies. For each section of text in the CORD19 JSON (titles, abstract parts and body parts), we injected a new block containing entities identified. The JSON block for each entity consisted of:

- a preferred name
- a unique identifier which in most cases was a URL for the relevant entity in its respective ontology
- a count of the number of times the entity was found in the section of text
- a list of the sentences in which the entity was found in the section of text
- a list of exact string locations for the sentences in which the entity was found in the section of text

By injecting these results back into the original JSON as an additional set of objects, we aimed to prevent any compatibility issues that would have arisen for individuals and groups who had already begun work with the initial release of this data.

To enable a more focused analysis, we also tabulated and transposed the annotated data (i.e. only sentences with annotations). In this case, we pivoted the data around the sentence, making the sentence the primary key, with attendant lists of entities identified in said sentence.

## Results

TERMite identified over 45 million entities within the initial CORD19 release, consisting of 62,746 unique identifiers. It should be noted that some terms are represented in multiple ontologies. This is particularly common with, for example, phenotypes from the Human Phenotype Ontology and indications from MeSH. (Table 6) summarises the annotations as a result of the NER process for each of the different vocabularies used. (Figure 1) gives an example of a sentence in which three annotations were found from three vocabularies.

**Figure 1.**
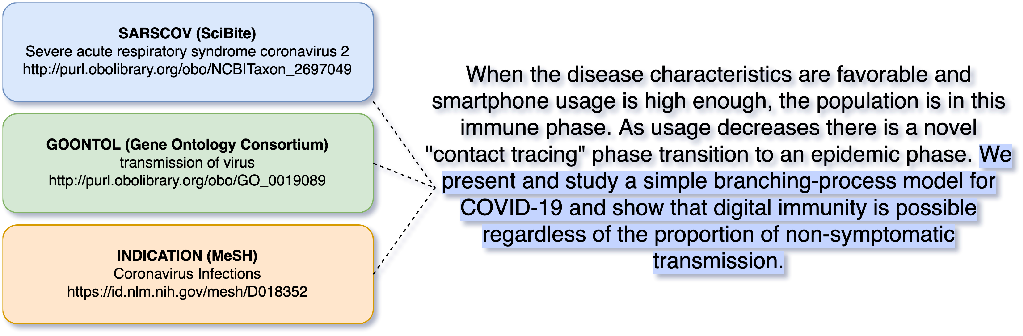
Example of a sentence containing three annotations. In this example, the virus name *COVID-19* is found, the GO term *transmission of virus*, and the MeSH indication *Coronavirus infection*.

The number of unique hits for each vocabulary type was many orders of magnitude less than the number of total hits found. This is very common in large, focused areas of research in this case anything COVID-19 relevant as the target virus will be mentioned many times in a single article.

Summaries of the most frequently found entities are shown in (Tables 3, 4 and 5). As can be observed in Table 3, many of the drugs found both in March and more recently have been heavily investigated by researchers around the world for their use as either a treatment (e.g. Hydroxychloroquine [8], Ribavirin [9]) or considering if they may play a role in poorer clinical outcome (e.g. Angiotensin [10]).

**Table 3.**
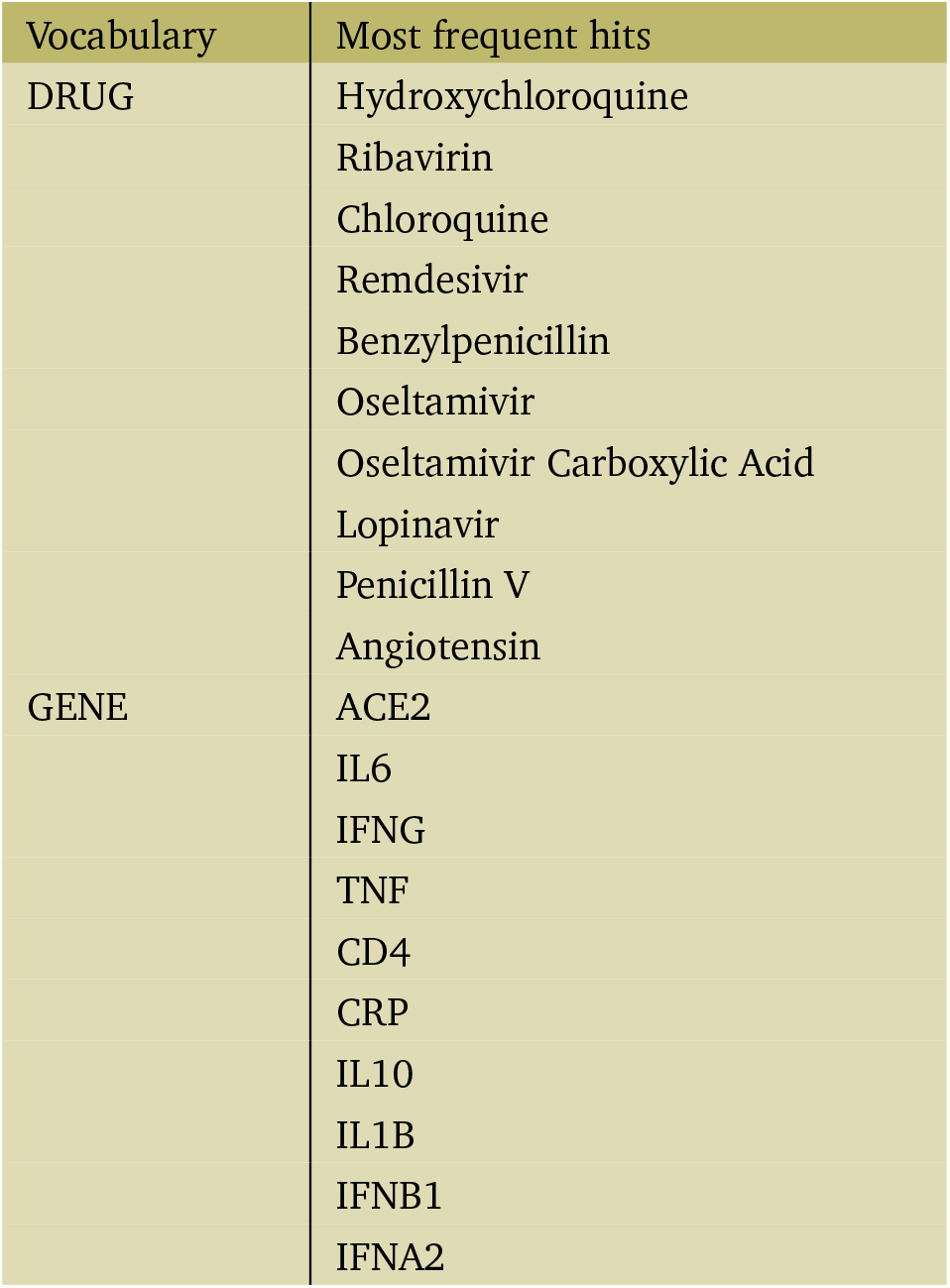
Most frequent annotations found for each vocabulary.

| Vocabulary | Most frequent hits |
| --- | --- |
| DRUG | Hydroxychloroquine |
|  | Ribavirin |
|  | Chloroquine |
|  | Remdesivir |
|  | Benzylpenicillin |
|  | Oseltamivir |
|  | Oseltamivir Carboxylic Acid |
|  | Lopinavir |
|  | Penicillin V |
|  | Angiotensin |
| GENE | ACE2 |
|  | IL6 |
|  | IFNG |
|  | TNF |
|  | CD4 |
|  | CRP |
|  | IL10 |
|  | IL1B |
|  | IFNB1 |
|  | IFNA2 |

**Table 4.** Most frequent annotations found for gene and drug vocabularies.

| Vocabulary | Most frequent hits |
| --- | --- |
| SARSCOV | Severe acute respiratory syndrome coronavirus 2<br>transmission of virus<br>SARS Coronavirus<br>viral life cycle<br>incubation period<br>viral genome<br>viral release from host cell<br>viral nucleocapsid<br>SARS coronavirus NS-1<br>viral strain |
| CORONAPROT | Angiotensin-converting enzyme 2<br>spike glycoprotein (SARS-CoV-2)<br>Spike glycoprotein<br>Replicase polyprotein 1ab<br>Replicase polyprotein 1a<br>nucleoprotein (SARS-CoV-2)<br>Nucleoprotein<br>3C-like proteinase (SARS-CoV-2)<br>Interleukin-6<br>RNA-directed RNA polymerase (SARS-CoV-2) |

**Table 5.** Most frequent annotations found for COVID19 specific vocabularies.

| Vocabulary | Most frequent hits |
| --- | --- |
| GOONTOL | binding<br>pathogenesis<br>behavior<br>growth<br>gene expression<br>signal transduction<br>cell death<br>immune response<br>learning<br>female pregnancy |
| HPO | pneumonia<br>fever<br>neoplasm<br>diarrhea<br>asthma<br>cough<br>sepsis<br>anxiety<br>depressivity<br>respiratory tract infection |

**Table 6.**
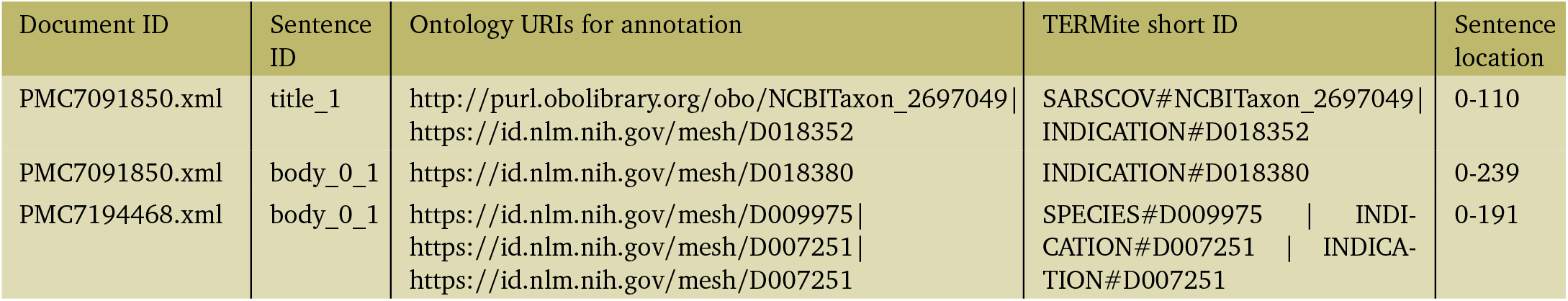
Example sentence level annotation.

| Document ID | Sentence ID | Ontology URIs for annotation | TERMite short ID | Sentence location |
| --- | --- | --- | --- | --- |
| PMC7091850.xml | title_1 | http://purl.obolibrary.org/obo/NCBITaxon_2697049 \| https://id.nlm.nih.gov/mesh/D018352 | SARSCOV#NCBITaxon_2697049 \| INDICATION#D018352 | 0-110 |
| PMC7091850.xml | body_0_1 | https://id.nlm.nih.gov/mesh/D018380 | INDICATION#D018380 | 0-239 |
| PMC7194468.xml | body_0_1 | https://id.nlm.nih.gov/mesh/D009975 \| https://id.nlm.nih.gov/mesh/D007251 \| https://id.nlm.nih.gov/mesh/D007251 | SPECIES#D009975 \| INDICATION#D007251 \| INDICATION#D007251 | 0-191 |

The ontology annotated data has now seen use in, for example, generating entity graphs using sentence cooccurrences as edges (Knowledge Graph Hub, 2020) or for combining evidence from multiple publication sources (Pub Annotation, 2020).

## Conclusions

The value of well-annotated and accessible data on a global scale has never been more acutely apparent. The CORD-19 dataset and call to action channeled this need for rapid and innovative action from the community to help research and development into tackling this global challenge. Within weeks of this call in March 2020, we have released to the community annotated sentences which mention entities of biomedical relevance to COVID19 and coronaviruses broadly, along with co-occurrences within sentences. The use of this data has already helped in developing knowledge graphs and data integration portals and it is our hope it will be put to further use in the data driven approaches used in 2020 and beyond.

## Acknowledgements

We would like to thank Chris Mungall for his suggestions on using the Protein Ontology and Lee Harland for his support on this work. We would also like to thank the Biohackathon community for their feedback on the data which has led to improvements in later versions.

